# The ultrastructure of *Shewanella oneidensis* MR-1 nanowires revealed by electron cryo-tomography

**DOI:** 10.1101/103242

**Authors:** Poorna Subramanian, Sahand Pirbadian, Mohamed Y. El-Naggar, Grant J. Jensen

## Abstract

Bacterial nanowires have garnered recent interest as a proposed **E**xtracellular **E**lectron **T**ransfer (EET) pathway that links the bacterial electron transport chain to solid-phase electron acceptors away from the cell. *In vivo* **f**luorescence **L**ight **M**icroscopy (fLM) imaging recently showed that *Shewanella oneidensis* MR-1 nanowires are extensions of the outer membrane that contain EET components. However, their fine structure and distribution of cytochrome electron carriers remained unclear, making it difficult to evaluate the electron transport mechanism along the nanowires. Here, we report high-resolution images of nanowires using **E**lectron **C**ryo-**T**omography (ECT). We developed a robust method for fLM imaging of nanowire growth on **e**lectron microscopy grids and used correlative light and electron microscopy to identify and image the same nanowires by ECT. Our results confirm that *S. oneidensis* nanowires are outer membrane extensions, and further reveal that nanowires are dynamic chains of interconnected **O**uter **M**embrane **V**esicles (OMVs) with variable dimensions, curvature, and extent of tubulation. Junction densities that potentially stabilize OMV chains are seen between neighboring vesicles in cryotomograms. Our ECT results also provide the first hints of the positions and packing of periplasmic and outer membrane proteins consistent with cytochromes. We observe tight packing of putative cytochromes along lateral patches that extend tens of nanometers, but not across the micrometer scale of whole nanowires. We therefore propose that electron transfer along nanowires involves a combination of direct hopping and diffusive events that link neighboring redox proteins.

## Introduction

Redox reactions are essential to all biological energy conversion strategies^1^. In respiratory organisms, free energy is harvested from the environment as electrons extracted from an electron donor (fuel) are transferred through the cellular electron transport chain to a terminal electron acceptor (oxidant). While most eukaryotes, including humans, are dependent on molecular oxygen (O_2_) as their terminal electron acceptor, anaerobic prokaryotes can acquire energy by employing a wide variety of alternative **E**lectron **A**cceptors (EAs). Like O_2_, many of these EAs can diffuse inside the cell, where they participate in redox reactions with intracellular electron transport chain components. However, **D**issimilatory **M**etal-**R**educing **B**acteria (DMRB) can also utilize insoluble EAs such as metal oxide minerals, to which the cellular membrane is impermeable, by transporting electrons across the cell envelope^2-5^. This **E**xtracellular **E**lectron **T**ransport (EET) process has important implications in renewable energy technologies, wastewater treatment, bioremediation, and global biogeochemical cycles^3,6-8^.

The Gram-negative bacterium *Shewanella oneidensis* MR-1 is the best-characterized DMRB model system^2,5,9,10^. Previous electrochemical, biochemical, genetic, and structural studies of *Shewanella* have identified an intricate network of redox proteins that traffic electrons from the inner membrane quinone pool through the periplasm and across the outer membrane^6,10,11^. A critical electron transfer module is the Mtr pathway, in which electrons are transferred from the periplasmic decaheme cytochrome MtrA to the outer membrane decaheme cytochrome MtrC through the transmembrane porin MtrB^12,13^. Under conditions of direct cell surface contact with minerals or electrodes, MtrC (and a partnering decaheme cytochrome OmcA) can transfer electrons directly to these solid EAs^14^. The EET rate from the surface-exposed cytochromes to such external surfaces can also be enhanced by interactions with secreted flavins that function either as cytochrome-bound cofactors^15-17^ or soluble shuttles capable of interacting with even more distant EA surfaces^18,19^. The *Shewanella* multiheme cytochromes MtrC and OmcA have also been found to localize along conductive outer membrane extensions known as bacterial nanowires, which are associated with **O**uter **M**embrane **V**esicles (OMVs)^20-22^. Nanowires may offer a pathway for extending the respiratory electron transport chain of cells up to micrometers away from the inner membrane, possibly even to other cells^22^.

Despite the potential impact in understanding cell-cell interactions, as well as for enhancing electron transfer at the biotic-abiotic interface in renewable energy technologies, we still lack a detailed understanding of the structure and electron transport mechanism of *S. oneidensis* nanowires. Physical measurements of conductance along the nanowires have revealed the importance of cytochromes as the charge carriers, but the measurements were made on dried samples so the distribution and conformation of the electron transport components may not be the same as *in vivo*^21^. Fluorescence-based *in vivo* techniques recently showed that *S. oneidensis* nanowires are extensions of the outer membrane and periplasm, and that nanowire production correlates with increased cellular respiration^22^. Additionally, immunolabeling experiments confirmed the localization of multiheme outer membrane cytochromes along the nanowire length^22^. However, the diffraction-limited resolution precluded visualization of the macromolecular details of the nanowire structure ^22^. Many other details remain unclear, including formation and stabilization mechanisms, cytochrome distribution, as well as the processes underlying the large morphological variation and dynamic nature of these filaments. One particular difficulty in previous studies is distinguishing nanowires from other filaments (flagella, pili, and dehydrated extracellular polymeric substances (EPS))^23,24^. **E**lectron **C**ryo-**T**omography (ECT), by contrast, can deliver high-resolution three-dimensional structural details of cellular structures. By capturing the specimen in a thin layer of vitreous ice, structures of interest are preserved in a fully-hydrated and essentially native state^25^.

Here we use ECT to capture the highest resolution images of bacterial nanowires to date. We have developed a novel experimental set-up allowing bacteria to form nanowires on an EM grid inside a perfusion flow imaging platform. Using fluorescent membrane staining, we monitored nanowire growth in real-time by live **f**luorescence **L**ight **M**icroscopy (fLM) and subsequently located and imaged the same nanowires by ECT. We discuss the challenges involved in retaining the fragile nanowire structures for EM imaging, and the methodology we developed to address these sample preparation issues. Our fLM and ECT results reveal the vesicular nature of *S.oneidensis* nanowires and shed light on a potential mechanism for their stabilization as OMV chains. The high resolution of ECT reveals the positions of the putative periplasmic and outer membrane multiheme cytochromes under near-native conditions. We discuss how these structural measurements inform and help refine proposed models^22,26,27^ for electron transport physics in *S. oneidensis* nanowires.

## Results

### Conditions for Reliable Nanowire Production for ECT

While OMVs and bacterial nanowires have previously been described in both planktonic and surface-attached *Shewanella* cultures using various methods such as EM, **A**tomic **F**orce **M**icroscopy (AFM), and fLM^20-22,28^, there has not been an extensive exploration of the optimal culturing and sample preparation workflows most suitable for detection of these structures. Here we utilized negative stain **T**ransmission **E**lectron **M**icroscopy (TEM) and ECT to assess both culturing and sample preparation steps that lead to robust formation and preservation of nanowires.

We first tested liquid cultures of *S. oneidensis* MR-1, either from continuous flow bioreactors (chemostats) operated under electron acceptor-limited conditions that trigger nanowire formation^20-22^, or from sealed serum bottles where *S. oneidensis* cells underwent a transition to electron acceptor limitation as they gradually consumed the available dissolved oxygen supplied from the headspace (Materials and Methods). Samples from both chemostats and serum bottles, either unfixed or fixed with glutaraldehyde, were visually assayed for nanowire formation by EM. Surprisingly, even in fixed cells, nanowires were rarely detected by either negative stain TEM or ECT, despite the presence of membrane blebs and OMVs (Figs. S1, S2). By contrast, similar to previous reports^23^, we observed an abundance of filaments by **S**canning **E**lectron **M**icroscopy (SEM) imaging of our liquid culture samples (data not shown). However, our investigations revealed that not all filaments observed in SEM could be identified as nanowires (additional experiments in progress), highlighting the need for improved methodology and care in interpretation of imaging results to distinguish nanowires from pili, flagella, and filamentous polymeric substances^24^.

Because nanowires in liquid cultures were only rarely observed by both ECT and negative stain TEM, we next tested surface-attached cultures. Building on our previous work utilizing surface-attached cultures to reveal the composition of *S. oneidensis* nanowires^22^, we developed a method for monitoring nanowire growth on EM grids inside a perfusion flow imaging platform by fLM (Fig.1). While nanowires were seen in this set-up by fLM, we observed only very few intact nanowires by either negative stain TEM or ECT, in unfixed or formaldehyde-fixed samples from this set-up (S3, S4). This suggests that nanowires are fragile structures. Fortunately, fixation with glutaraldehyde stabilized the nanowires, enabling us to reliably visualize the structures by **C**orrelative **L**ight and **E**lectron **M**icroscopy (CLEM) (Fig. S5, Movies S1 and S2). We conclude that nanowires are more frequent and consistently present in surface-attached cultures, but comparatively uncommon in liquid cultures under our experimental conditions.

**Fig. 1:**
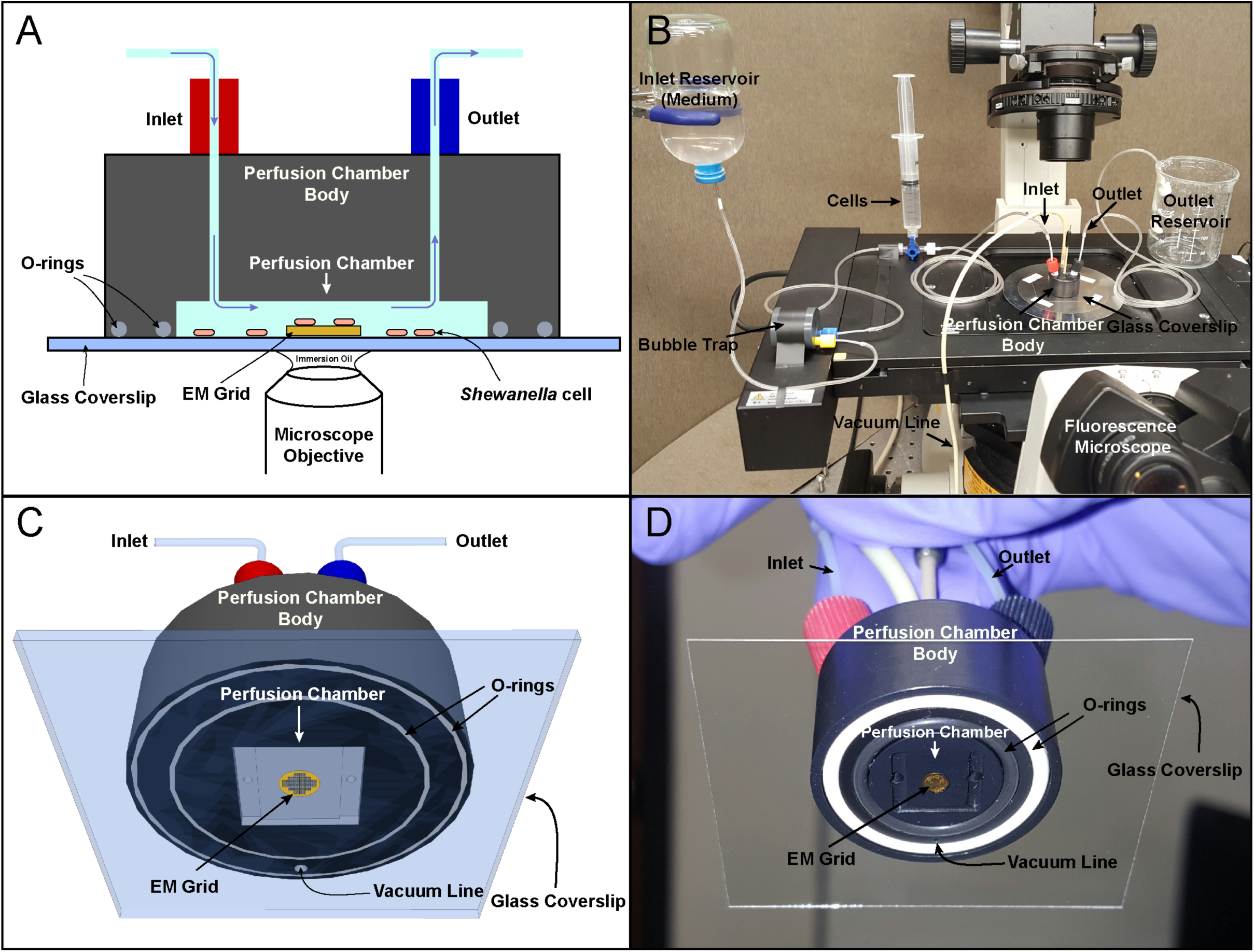
Schematic and actual images of the perfusion flow imaging platform (objects not drawn to scale). (A) Cross-sectional and (B) Three-dimensional (3D) views of the perfusion flow imaging platform. An electron microscopy (EM) grid is glued to a glass coverslip that seals the perfusion chamber. S. oneidensis cells injected into the sealed chamber attach to the grid surface and are sustained by a continuous flow of the medium. Cells are labeled with the fluorescent membrane dye FM 4-64FX and monitored real-time for nanowire growth using an inverted fluorescent microscope placed under the perfusion chamber. (C) 3D schematic and (D) Image of the perfusion chamber interior with an attached EM grid.

### Live Fluorescence Microscopy of Nanowire Growth on EM Grids

Our optimized perfusion flow imaging platform set-up consists of a microliter-volume laminar perfusion flow chamber placed on an inverted fluorescence microscope, where an EM grid-attached glass coverslip seals the chamber (Fig. 1). *S. oneidensis* cells are then introduced into the chamber, where they attach to the surface of the EM grid, and sterile media is flowed into the chamber throughout the experiment. The laminar Poiseuille flow (no mixing from upper layers), combined with negligible flow speed at the surface-solution interface (no-slip condition), where the cells are located, results in O_2_-limiting conditions that trigger nanowire formation directly on the EM grid^22^.

Using this set-up, we observed the formation of nanowires live on the EM grid surface with the fluorescent membrane dye FM 4-64FX. Cells were located relative to grid holes by fLM (Fig. 2 and Movie S3) to allow registration with subsequent EM imaging. It has been shown previously that the conductance of nanowires depends on the presence of the outer membrane multiheme cytochromes MtrC and OmcA^21^, and these cytochromes have been detected using immunofluorescence along membrane-stained nanowires from perfusion flow cultures^22^, so the nanowires we observe are likely conductive.

**Fig. 2:**
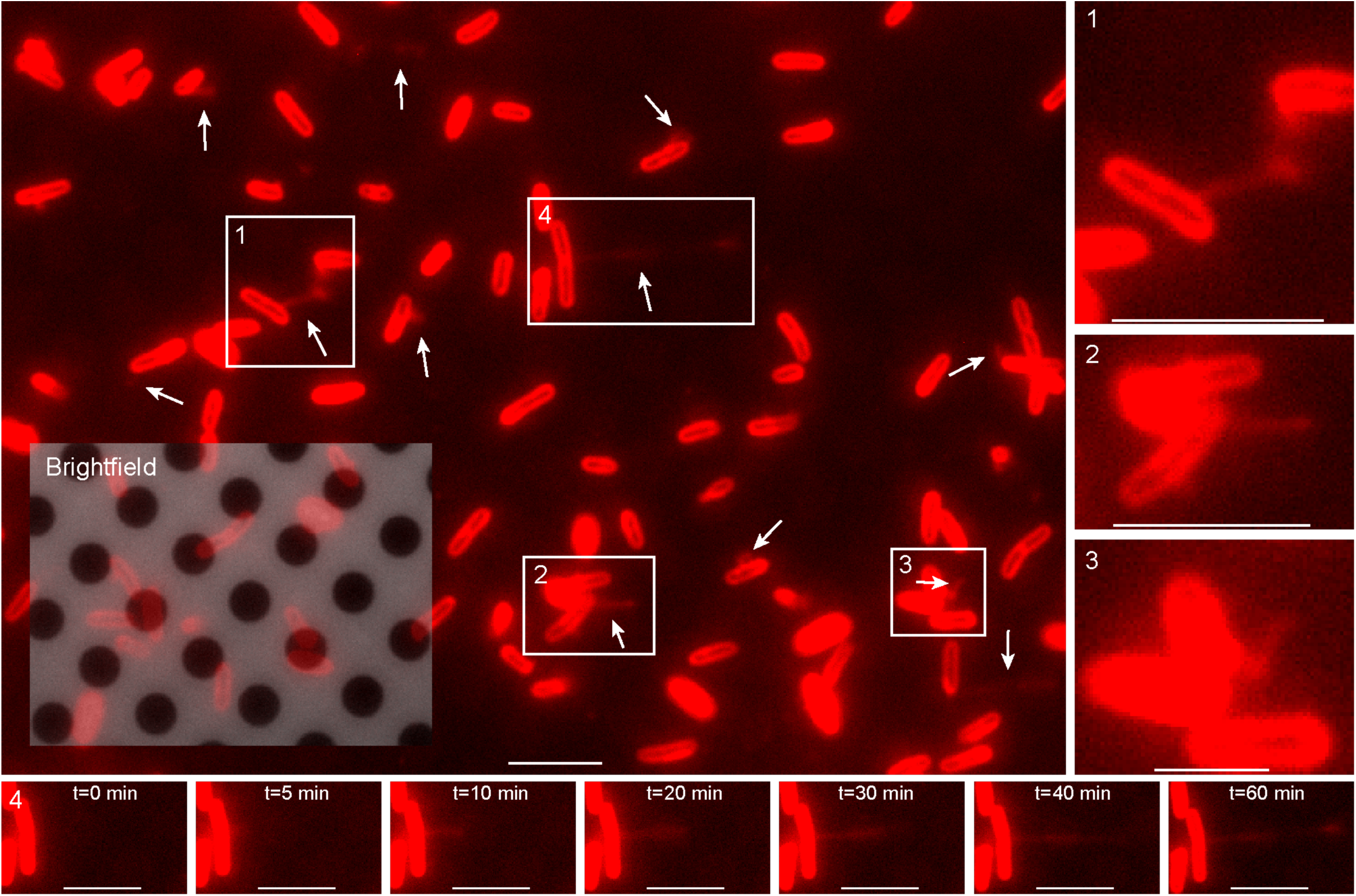
Live *in vivo* observation of the formation of *S. oneidensis* nanowires (white arrows) on an EM grid (Scale bar: 5 µm). Inset is an overlap of red fluorescence and reflective brightfield channels, revealing both the holey carbon film coating the EM grid and the fluorescently labeled cells attached to it. Movie S3 is a time-lapse movie of this figure. (1, 2, and 3) Enlarged views of boxed regions from the main panel. (Scale bars in 1, 2, and 3: 5, 5, and 2 µm, respectively). (4) Time-lapse images of the growth of a single nanowire from boxed region 4 in the main panel. *t=0* min is an arbitrary starting timepoint. (Scale bars: 5 µm).

### Nanowires are Dynamic Chains of Interconnected OMVs Stabilized by Junction Densities

For ECT, grids from the perfusion flow imaging platform were removed, plunge-frozen and transferred to the electron microscope, where the fLM-identified nanowires were located and imaged (Fig. 3). ECT images confirmed that nanowires are outer membrane extensions, with the two leaflets of the lipid bilayer clearly resolved along their length (Fig. 4A-B). Cryotomograms revealed nanowires to be chains of interconnected OMVs in both unfixed (Fig. 4C) and fixed samples (Fig. 4D-G). Previous fLM and AFM work showed that nanowires cover a range of morphologies from apparently smooth tubular extensions to clearly distinguishable OMV chains^22^. Here, we observed that, with the exception of one smooth filament (Fig. S6), all nanowires including those that appeared smooth in fLM were, at high resolution, distinguishable as OMV chains (Figs. 3 and 4). The images also captured vesicle budding (Fig. 4B), a process that underlies the initial stage of OMV production^29^. Importantly, ECT also allowed us to clearly distinguish between pili, flagella and nanowires – the three known extracellular appendages in *S. oneidensis* (Fig. 4D-G, Movies S4 and S5).

**Fig. 3:**
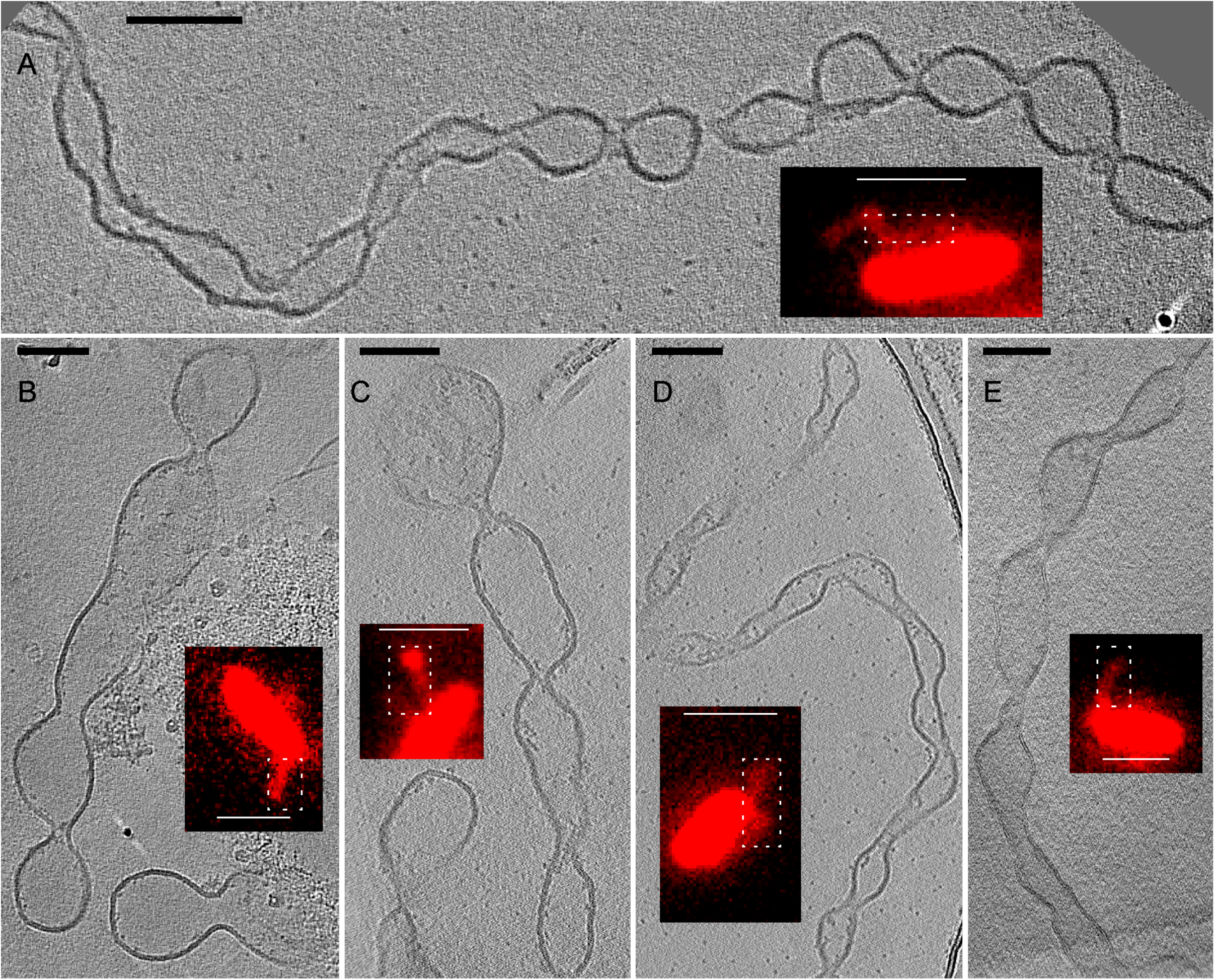
Targeting dynamic nanowire structures of*S. oneidensis* for ECT using **C**orrelative **L**ight and **E**lectron **M**icroscopy (CLEM). To visualize the structureof nanowires by ECT, target locations on fixed and plunge-frozen EM grids, from the perfusion flow imaging platform, were imaged, revealing the OMV chain morphology of the nanowires. (A, B, C, D, and E) Representative images from ECT, with corresponding fluorescent light microscopy image insets. (ECT scale bars: 100 nm, fLM scale bars: 2 µm). White dotted boxes in the fLM images indicate the corresponding approximate regions imaged in ECT. The ECT images shown are tomographic slices from three-dimensional reconstructions. See also Fig. S5 and Movies S1 and S2.

**Fig. 4:**
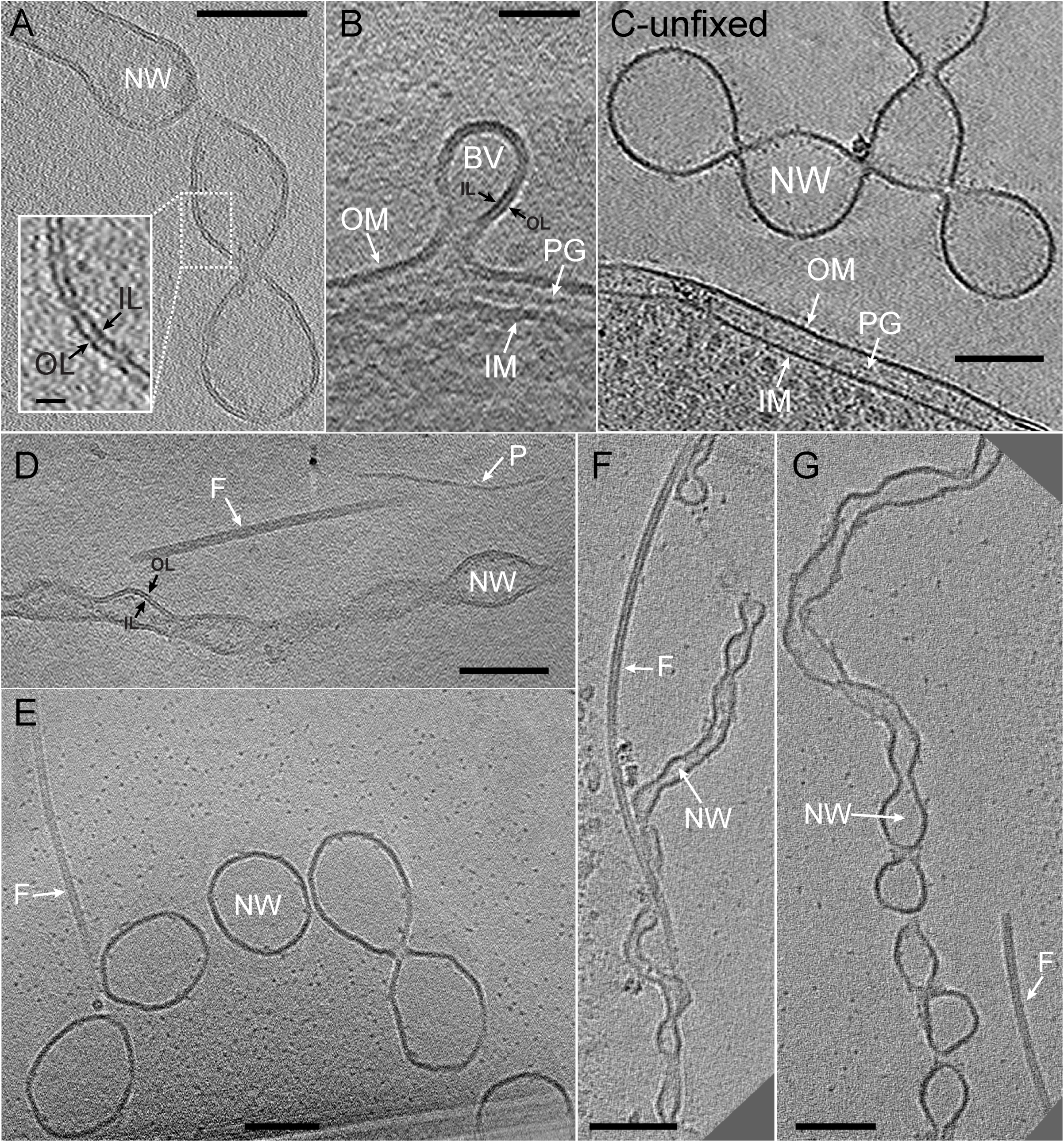
Electron cryo-tomography (ECT) images of *S. oneidensis* nanowires.(A) Nanowire membrane bilayer is clearly resolved. Inset is enlarged view of boxed region with the inner and outer leaflets indicated with arrows. (Scale bar: 100 nm, inset scale bar: 10 nm). (B) A budding vesicle (BV) emerging as an extension of the cellular outer membrane. A similar process perhaps underlies the initial stages of nanowire formation. (Scale bar: 50 nm). (C) Nanowire from an unfixed chemostat sample exhibits identically branched OMV chain morphology as observed in both unfixed and fixed samples from the perfusion flow imaging platform. (Scale bar: 100 nm). See also Figs. S4, S9 and Movie S12. (D) A nanowire, a flagellum and a pilus next to each other allowing direct comparison of their sizes and morphologies, indicating that ECT facilitates the identification and distinguishability of different extracellular appendages in *S. oneidensis*. (Scale bar: 100 nm). See also Movie S4. (E, F, and G) ECT reveals nanowires are of varying thicknesses and degrees of tubulation. Next to each nanowire is a flagellum that can act as a molecular marker for comparison of varying nanowire dimensions. (Scale bars: 100 nm). See also Movie S5 corresponding to (F). NW-nanowire, Fflagellum, P-pilus, BV-budding vesicle, OM-outer membrane, IM-inner membrane, PG-peptidoglycan, OL-outer leaflet, IL-inner leaflet

Electron-dense regions were observed at the junctions connecting neighboring vesicles throughout the length of the nanowires in both fixed and unfixed samples (Figs. 5A, S7 and Movie S6). This finding points to yet unknown molecules that potentially facilitate the constriction of the membrane to allow OMV connections, and is consistent with the fLM observations of the nanowires as dynamic structures capable of growth, retraction, and reversible transition between OMV chain and individual vesicle morphologies (Figs. 5B, 5C, and Movies S7, S8, S9, S10).

**Fig. 5:**
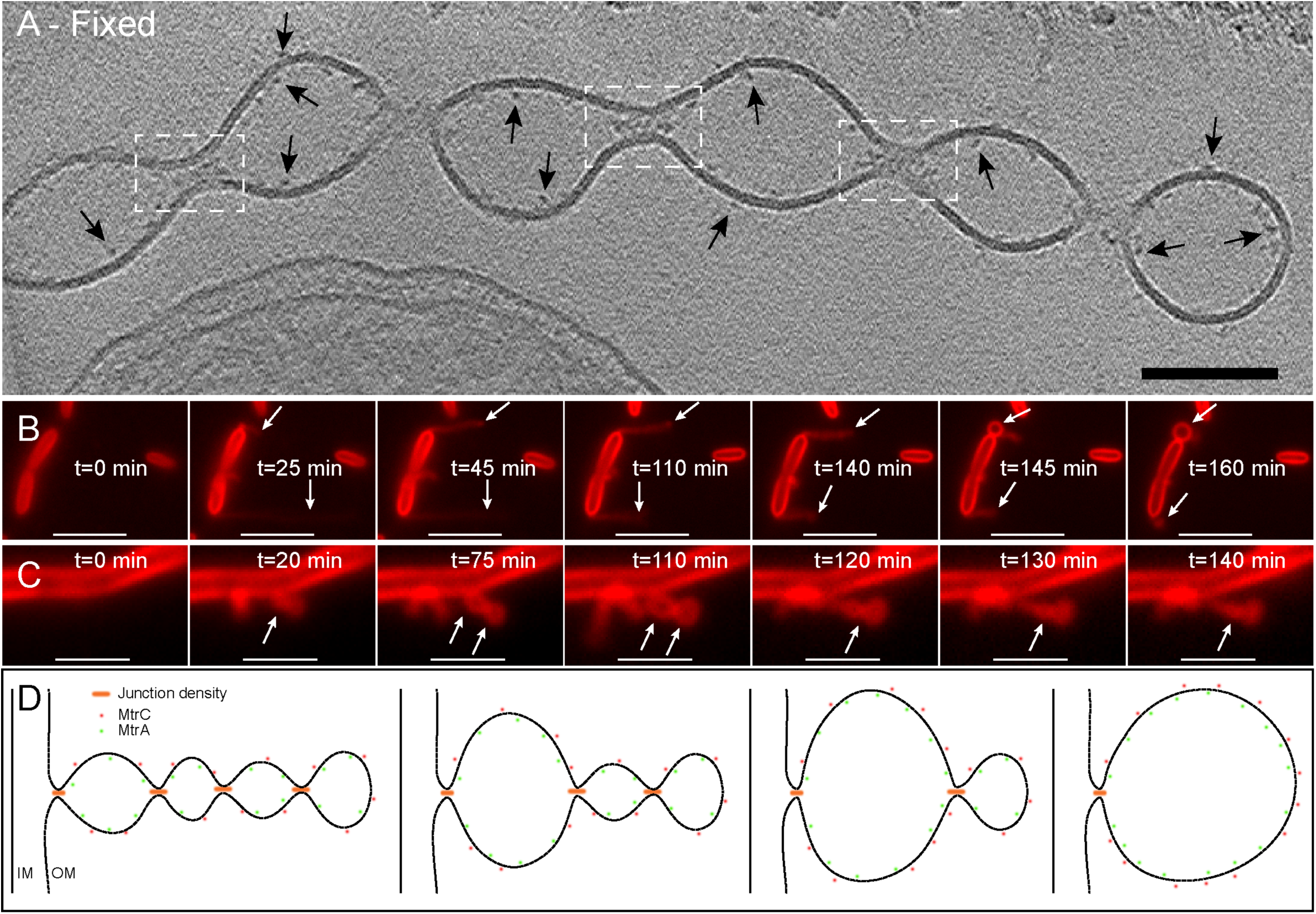
Proposed model for the formation and stabilization of OMV chains. (A)ECT image of a chemically fixed nanowire reveals the presence of densities at junctions that connect one vesicle to the next along the OMV chain (white dashed boxes). All densities are not visible in the tomographic slice in (A), while Movie S6 is a 3D reconstruction revealing the densities present at every junction. In addition, densities possibly related to decaheme cytochromes can be observed on the interior and exterior of the outer membrane along the nanowire (arrows). (Scale bar: 100 nm). See also Fig. S7. (B and C) Time-lapse fluorescence images recorded real-time in the perfusion flow imaging platform monitoring the growth and transformation of a nanowire from (B) An apparently long filament (OMV chain morphology) to a single large vesicle (indicated by arrows) in *S. oneidensis Δflg* (a mutant strain lacking flagellin genes). (See also Movie S7). (Note that nanowires from wild-type cells also exhibit a similar behavior as shown in Movie S8) and (C) A large vesicular morphology to an apparently smoother filament (OMV chain morphology) (indicated by arrows) in wild-type *S. oneidensis* MR-1 cells. See also Movies S9 and S10. The cells and the nanowires in (B) and (C) are stained by the membrane stain FM 4-64FX. (Scale bars in B and C: 5 and 2 µm, respectively). (D) Schematic depicting a hypothesis for the formation and stabilization mechanism of OMV chains: Junction densities on the interior of the nanowire facilitate the constriction of the membrane, enabling the formation of an OMV chain. These constriction densities can be removed or added to facilitate transformation of an OMV chain to a large vesicle or vice versa as observed in (B) and (C) respectively.

### ECT Reveals the Distribution of Periplasmic and Outer Membrane Proteins Consistent with Multiheme Cytochromes

The outer membrane cytochromes MtrC and OmcA have been shown to localize along the nanowire length^22^ and are essential for solid-state conductance of nanowires in *S. oneidensis*^21,22^. The packing density of these cytochromes is crucial in determining the mechanism of electron transport along nanowires, but has remained unknown. Here, using ECT, we observe electron-dense particles on the interior and exterior of the nanowires’ membrane consistent with periplasmic and outer membrane cytochromes (Fig. 6A). Fig. 6B overlays available structures of the decaheme cytochromes MtrA^30^ and MtrC^16^ on representative interior and exterior densities, respectively, highlighting the similarity in overall shape and size of these structures to the observed EM densities. From the cryotomograms, it is possible to represent all the observed interior and exterior densities as model points and reconstruct 3-dimensional isosurfaces that represent both the nanowire structure and the putative EET proteins (Fig. 6C and Movie S11). While we observed sections up to ~70 nm and ~75 nm where the exterior and interior densities clustered closely with 7.3 nm (SD=2.1 nm) and 8.9 nm (SD=2.0 nm) center-to-center distances, respectively (Fig. 6D-E), we did not observe a continuous crystalline-like packing of these densities along the entire nanowire length (Fig. 6H). Instead, the outer membrane and periplasmic densities were distributed over a range of center-to-center spacings, from 4.9 to 32.5 nm and 5.0 to 29.0 nm, respectively (Fig. 6I).

**Fig. 6:**
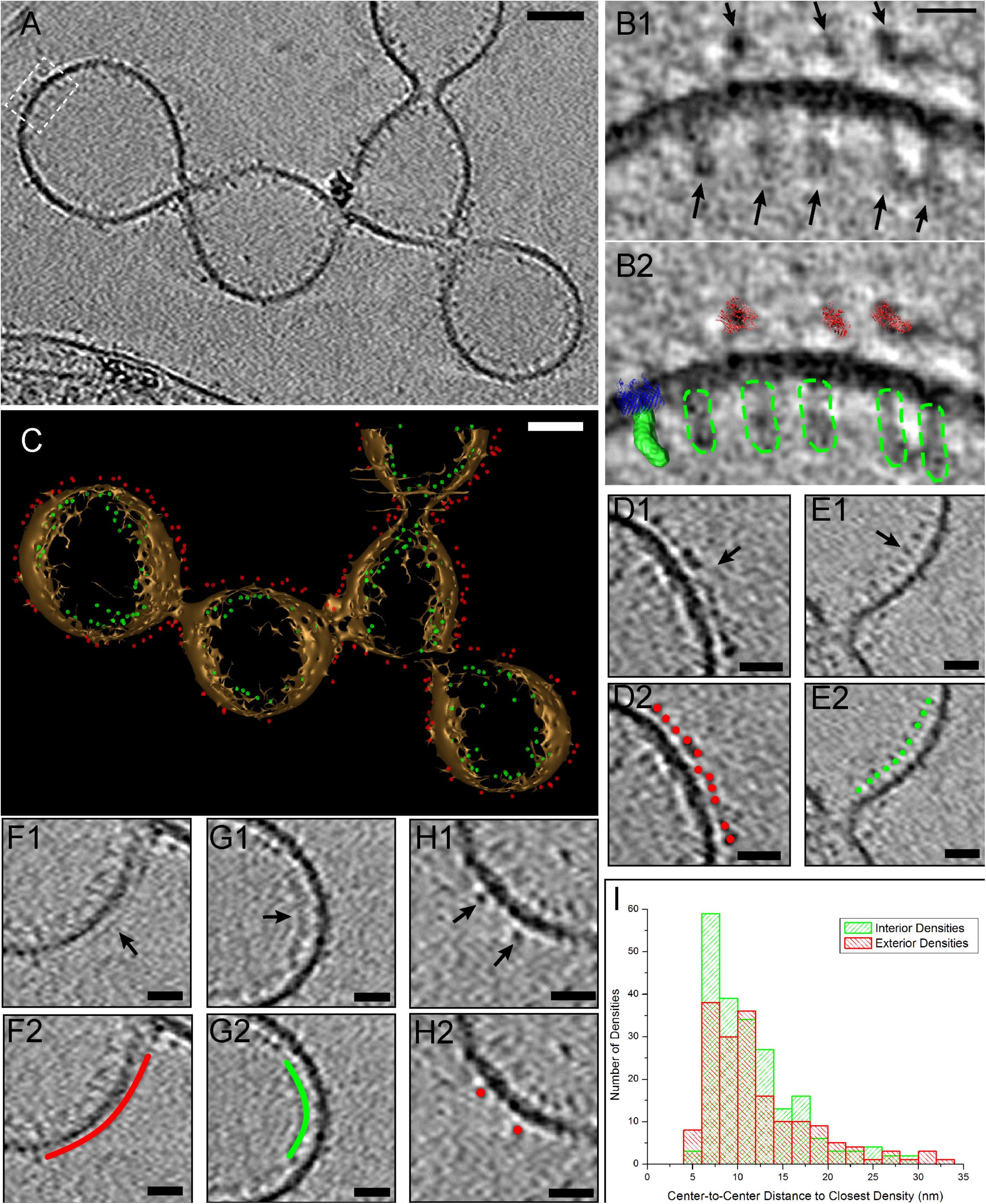
Positions and packing of decaheme cytochromes along the nanowire length in *S. oneidensis* (A) ECT image of an unfixed nanowire showing densities on both the interior and exterior of the outer membrane corresponding to putative MtrA and MtrC cytochromes respectively (Scale bar: 50 nm). (B1) Enlarged view of boxed area from (A). (B2) Comparison of EM densities in (B1) with the crystal structure of MtrC, low resolution SAXS model of MtrA, and the MtrB homolog LptD (only the LptD structure was used for this model from the LptD-LptE two-protein crystal structure), highlighting the similarity in overall shape and size of these structures to the observed EM densities. Red: MtrC crystal structure, Green: surface view of MtrA SAXS model, Blue: LptD crystal structure, Dotted green: outline of putative MtrA densities on the EM map. (Scale bar: 10 nm). (C) 3D isosurface view of the nanowire in (A) with all the interior and exterior densities (putative MtrA and MtrC respectively) represented as model points in green and red respectively. (Scale bar: 50 nm). See also Movie S11. (D-G) Representative regions from the nanowire in (A) demonstrating differences in packing density of MtrA and MtrC. (D1) Relatively closely packed exterior densities of putative MtrC with an average center-to-center inter-density distance of 7.3 nm (SD=2.1 nm). (E1) Relatively closely packed interior densities of putative MtrA with an average center-to-center inter-density distance of 8.9 nm (SD=2.0 nm). Continuous (F1) interior and (G1) exterior densities that may be related to tightly packed MtrA and MtrC respectively. (H1) Isolated exterior densities of putative MtrC. (D2), (E2), (F2), (G2), and (H2) are duplicates of (D1), (E1), (F1), (G1), and (H1) respectively, with model points or lines highlighting the interior (green) and exterior (red) densities. (I) Histogram showing distribution of center-to-center distances of all putative MtrAs (in green) and MtrCs (in red).

## Discussion

Here we show high-resolution images of nanowires in *S. oneidensis* using electron cryo-tomography. We found nanowires to be OMV chains possibly stabilized by constriction densities at the junctions. Bacterial membrane extensions have been reported in multiple organisms: ‘Nanopods’ in *Comamonadaceae* including *Delftia*^31^, ‘outer membrane tubes’ in *Francisella novicida*^32^, ‘periplasmic tubules’ in *Chlorochromatium aggregatum*^33^*, ‘* membrane tubules’ in *Salmonella typhimurium*^34^, ‘nanotubes’ connecting *Escherichia coli* cells to each other and to *Acinetobacter baylyi* cells^35^, and ‘connecting structures’ that allow exchange of material between *Clostridium acetobutylicum* and *Desulfovibrio vulgaris* cells^36^. However, membrane extensions in the form of OMV chains have only recently been discovered and much remains unknown about their formation mechanism and specific function^37^. In the Gram-negative *Shewanella vesiculosa*^38^ and *Myxococcus xanthus*^39,40^ and the Gram-positive *Bacillus subtilis*^41^, membrane extensions in the form of OMV chains, similar to those reported here, have been observed using cryo-EM with implications for cell-cell connections in the latter two examples. While the *S. oneidensis* nanowires are thought to function as electron conduits^21^, their structural similarity to these previous reports highlights the significance of imaging nanowires as a model system to study the formation of OMV chains.

In order to find a condition that consistently and frequently produced intact nanowires for ECT imaging, we systematically tested different methods of growth and sample preparation conditions, as summarized in Fig S8. Using surface-attached cultures fixed with glutaraldehyde stabilized the nanowires, enabling us to record high-resolution cryotomograms of a number of *S. oneidensis* nanowires as shown in Fig 4. The OMV chain morphology exhibited by these nanowires is unlikely to be an artifact of fixation since we also observed a similar OMV chain architecture in nanowires from unfixed samples (Figs 6A, S4). While flagella and pili were identified as smooth filaments that measured ~10 and ~3 nm in thickness, respectively, nanowires varied in thickness typically from ~20 to 170 nm (Fig. 4D-G), depending on the size and extent of tubulation of the constituent OMVs. This is in contrast to previously reported AFM measurements of ~10 nm for air-dried nanowires. This discrepancy arises from the difference in sample preparation between the two methods. While in AFM dehydration causes nanowires to collapse to an ~10 nm thickness, roughly corresponding to two lipid bilayers, ECT preserves samples in a hydrated near-native state, leading to more accurate estimates of nanowire thickness. An interesting feature we observed is the ability of the vesicle chains to branch (Figs. 6A, S9 and Movie S12), which may offer the advantage of increasing the likelihood of contacting terminal solid-phase electron acceptors in the environment. To our knowledge this is the first report of branching observed in bacterial membrane extensions. Nanowires were also found to be flexible (Fig S10), potentially improving their ability to contact solid-phase EAs.

Our ECT images of *S. oneidensis* nanowires reveal that individual vesicles open into each other, share a continuous lumen, and thus form a chain of vesicles that are internally connected. This OMV architecture is reminiscent of the “pearls on a string” morphology caused by the pearling instability that transforms membrane tubes into a string of interconnected vesicles^42,43^. The transformation may be caused by an increase in membrane tension that can be stimulated in multiple ways, including osmotic gradient^44^, mechanical perturbation^42,43^, elongational flow^45^, electric field^46^, bilayer asymmetry^47^, nanoparticle adsorption onto the inner leaflet^48^, or polymer anchorage onto a membrane^49,50^. Our observation of densities at the junctions of neighboring vesicles in both fixed (Fig. 5A and Movie S6) and unfixed (Fig. S7) nanowires is consistent with the latter mechanism of polymer anchorage onto a membrane in which ‘constriction densities’ or ‘junction densities’ interact with the nanowire membrane, resulting in the formation of the OMV morphology (Fig 5A arrows). As schematized in Fig. 5D, addition and removal of such constriction densities may also explain the dynamic behavior exhibited by nanowires, where long nanowires transition to and from individual vesicles (Fig. 5B-C, Movies S7, S8, S9, S10).

The solid-state conductance of *S. oneidensis* nanowires has been linked to the presence of the outer membrane decaheme cytochromes MtrC and OmcA^21^. These cytochromes are localized along the length of *S. oneidensis* nanowires and are thought to mediate electron transport by a multistep redox hopping mechanism^22,26^. While the intra-protein hemes’ arrangement within MtrC and OmcA allows sequential tunneling (hopping) through the heme chain^51,52^, the packing density and orientation of these cytochromes are critical parameters that determine the mechanism of putative inter-protein electron transfer along the entire nanowire. However, prior to this work, little was known about the packing density of MtrC and OmcA molecules along nanowires.

The nanowires in ECT showed densities both on the inside and outside of the membrane (Fig. 6A), features consistent with the periplasmic and outer membrane proteins, respectively, and as expected from immunofluorescence measurements^22^. To determine whether the outside densities match the size of MtrC, we overlaid the crystal structure of MtrC^16^ onto three of these densities, as illustrated in Figure 6B. Since these densities did not appear symmetric on the EM map, and since the orientation of MtrC at the cellular outer membrane is unknown, we overlaid the MtrC crystal structure in the orientation that best matched each density. Using this approach, the size of the outer membrane features was found to be consistent with MtrC. This approach cannot distinguish between MtrC and other *Shewanella* outer membrane decaheme cytochromes, such as MtrF and OmcA, all of which have substantial structural homology^16^. The size of these proteins is at the resolution limit of tomography and additional experiments are needed to obtain further details about the interaction and association of MtrC with other proteins in the MtrCAB complex^13,53^ that is proposed to form a contiguous EET conduit from the periplasm to the cellular exterior. We applied a similar approach to compare the interior nanowire densities with the periplasmic decaheme cytochrome MtrA. The interior densities were more oblong than their outside counterparts, an observation consistent with the rod-like shape of MtrA previously revealed by **S**mall-**A**ngle **X**-ray **S**cattering (SAXS)^30^. By overlaying this low-resolution SAXS model on the EM map, the internal densities were found to be consistent in size and shape with MtrA (Fig. 6B). While the structure of MtrB is not yet known, we overlaid the crystal structure of a similarly-sized protein (LptD from *Salmonella enterica*^54,55^) in Figure 6B, and found that the size of the porin matches the width of the bilayer as expected. Taken collectively, our analyses highlight the similarity in overall shape and size between multiheme cytochromes and the observed EM densities.

The isosurface representation of the nanowires, including the placement of the detected periplasmic and outer membrane proteins (Fig. 6C and Movie S11), allows a holistic evaluation of different inter-protein electron transfer mechanisms. Remarkably, we observed outer membrane (Fig. 6D) and periplasmic (Fig. 6E) proteins clustering closely over segments of the nanowire. The center-to-center distances between neighboring proteins within the tightest-packed segments were 7.3 nm (SD=2.1 nm) and 8.9 nm (SD=2.0 nm) for the outer membrane and periplasmic proteins, respectively. Taking the overall dimensions of MtrC^16^ (~9×6×4 nm) and the locations of the hemes (including terminal hemes at the protein edges) into account^16^, the center-to-center distances point to the possibility of direct electron tunneling (requiring < 2 nm separation^1^) between terminal hemes of neighboring outer membrane cytochromes within segments. However, such a crystalline-like packing of cytochromes was not observed over the micrometer lengths of whole wires (Fig. 6). Instead, we observed a wide distribution of center-to-center spacings, presented for both the periplasmic and outer membrane densities in Fig. 6I. Since center-to-center spacings beyond 11 nm and 7 nm for MtrC and MtrA, respectively, do not allow direct electron transfer between neighboring cytochromes (see Materials and Methods), intermediate diffusive events are required to link the hemes of neighboring proteins beyond such distances.

This may be accomplished by lateral diffusion of the multiheme cytochromes, resulting in collisions and electron exchange between neighboring cytochromes. For example, Paquette *et al.*^56^ suggested that OmcA, which interacts with MtrC and is attached only by a lipidated cysteine at the N-terminus, is mobile on the surface of *Shewanella*. Using 3 μm^2^/s as a representative value for the diffusion coefficient of integral membrane proteins of similar size^57^, the median inter-cytochrome distance can be traversed in 10^-5^ s (Table S1, Materials and Methods), a time scale comparable with the electron residence time in the heme chains of the individual cytochromes, estimated from calculated and measured electron flux through MtrF (10^4^ s^-1^)^51,52^ and MtrCAB^14^. In addition, the electron transfer rate can be enhanced by diffusion of redox active molecules between cytochromes. In this context, it is important to note that the *Shewanella* decaheme cytochromes have flavin binding sites^16^, and flavins are known to enhance EET^17^. The precise values of the diffusion coefficients (MtrC/OmcA proteins in the membrane, the likely faster MtrA diffusion within the periplasm, and the effect of crowding) are unknown. The preceding analysis is therefore intended for heuristic reasons, and to motivate future studies targeting the diffusive dynamics of electron carriers in bacterial nanowires. In summary, our ECT imaging revealed particles consistent in size and morphology with decaheme cytochromes and their distribution along nanowires. Although we do not yet know whether all of the densities observed on the inside and the outside of the membrane correspond to MtrA and MtrC respectively (since we cannot distinguish between MtrC and other structural homologs, such as MtrF and OmcA, and there may be other membrane proteins as well), it is already clear that cytochromes are not tightly packed along the entire length of the nanowire. This irregular packing of cytochromes means that EET along whole nanowires likely involves a combination of direct electron transfer within segments of closely packed cytochromes and cytochrome diffusion to bridge larger length scales.

## Acknowledgements

We thank Drs. Yi-Wei Chang and Matthew Swulius for help preparing Figures 6B and 6C, respectively. We are grateful to Dr. Sean J. Elliott for providing the SAXS model file for MtrA^30^ used in Figure 6B. Thanks to Dr. Catherine Oikonomou for helping edit the manuscript. P.S. acknowledges support by the Caltech **C**enter for **E**nvironmental **M**icrobial **I**nteractions (CEMI). Work in the laboratory of G.J.J. is supported by the Howard Hughes Medical Institute. The *in vivo* nanowire imaging platform and mapping of EET proteins is funded by Air Force Office of Scientific Research PECASE award FA955014-1-0294 to M.Y.E-N. Modeling of ET kinetics and partial support for S.P. are funded by the Division of Chemical Sciences, Geosciences, and Biosciences, Office of Basic Energy Sciences of the US Department of Energy through grant DE-FG02-13ER16415 to M.Y.E-N.

## Materials and Methods

### Chemostat Growth Conditions

*S. oneidensis* MR-1 cells were grown in continuous flow bioreactors (BioFlo 110; New Brunswick Scientific) with an operating liquid volume of 1 L, as previously described^20-22^. 5 mL of a stationary-phase overnight Luria-Bertani (LB) culture was injected into the bioreactor, while **D**issolved **O**xygen **T**ension (DOT) was maintained at 20% by adjusting the ratio of N_2_/air mixture entering the reactor (using automatic mode). After 20 hours, and as the culture reached stationary phase, continuous flow of the medium was started with a dilution rate of 0.05 h^-1^ while DOT was still maintained at 20%. After 48 hours of aerobic growth under continuous flow conditions, the DOT was manually reduced to 0% by adjusting the N_2_/air mixture entering the reactor. O_2_ served as the sole terminal electron acceptor throughout the experiment. pH was maintained at 7.0, temperature at 30 °C and agitation at 200rpm to minimize mechanical shear forces. 40 hours after DOT reached 0%, samples were taken from the chemostat for TEM imaging.

### Serum Bottle Growth Conditions

*S. oneidensis* MR-1 was grown overnight in Luria-Bertani (LB) broth at 30 °C up to an OD_600_ of 2.4-2.8. 200 µL of this overnight culture was added to each of two duplicate, sealed, and autoclaved 100-mL serum bottles containing 60 mL of a defined medium^22^. One of the two bottles acted as a control and was not used for imaging. To the control bottle, 5µM resazurin was added to indicate the O_2_ levels in the medium. The bottles were then placed in an incubator at 30 °C, shaking at 150 rpm until the color due to resazurin in the control bottle completely faded, indicating anaerobic conditions. We then took 200 µL of sample from the bottle that did not contain resazurin for TEM imaging.

### Perfusion Flow Imaging Platform

The perfusion flow imaging platform was used as described previously^22^, with some modifications. *S. oneidensis* MR-1 was grown overnight in Luria-Bertani (LB) broth at 30 °C up to an OD_600_ of 2.4-2.8 and washed twice in a defined medium^22^. A glow-discharged, X-thick carbon-coated, R2/2, Au NH2 London finder Quantifoil EM grid (Quantifoil Micro Tools) was glued to a 43 mm × 50 mm No. 1 glass coverslip using waterproof silicone glue (General Electric Company), applied to two opposite edges of the grid, and let dry for ~30 min. Using a vacuum line, the perfusion chamber (C&L Instruments, model VC-LFR-25) was sealed against the grid-attached glass coverslip and placed on an inverted microscope (Nikon Eclipse Ti-E) that continually imaged the grid surface. ~10 ml of the washed culture was injected into the chamber slowly to allow cells to settle on the grid surface, followed by flow of sterile defined medium from an inverted serum bottle through a bubble trap (Omnifit, model 006BT-HF) into the perfusion chamber inlet. The serum bottle was pressurized by N_2_ in the headspace to sustain a flow rate of 5±1 µL/s. After ~2 hrs of perfusion flow, cells on the grid surface began to produce nanowires. Cells and nanowires were visualized by the fluorescent membrane stain FM 4-64FX that was present in the flow medium throughout the experiment (25 µg in 100 mL of medium). Subsequently, the flow of medium was stopped and the perfusion chamber was opened under sterile medium. If fixed, the sample (cells on EM grid-attached coverslip) was treated with either 2.5% glutaraldehyde for 15 min or 4% formaldehyde for 60 min. The grid was then detached from the coverslip by scraping off the silicone glue at the grid edges using a 22G needle, and rinsed by transferring 3 times in deionized water, before using for TEM imaging.

### Negative Stain Transmission Electron Microscopy (TEM)

#### (i) Chemostat

Samples taken from the chemostat were immediately fixed with 2.5% glutaraldehyde and stored at 4 °C. 2ul of this fixed sample was spotted on a glow-discharged, X-thick carbon-coated, R2/2, Au NH2 London finder Quantifoil EM grids (Quantifoil Micro Tools) and let sit for 2 min. Any remaining liquid was blotted off gently using a kimwipe and the grid was stained with 1% uranyl acetate for 2 min before gently blotting the remaining stain. The grid was then allowed to dry for a day at room temperature before imaging in TEM.

#### (ii) Perfusion Flow Imaging Platform

The fixed grid from the perfusion flow imaging platform was blotted to remove excess liquid, and stained with 1% uranyl acetate for 2 min before gently blotting the remaining stain. To obtain an fLM image for correlation with EM, the grid was rehydrated with deionized water and reimaged on the inverted fluorescent microscope. The grid was then blotted again and let dry at room temperature before imaging in TEM.

### Electron Cryo-Tomography (ECT)

ECT samples were prepared as described previously^58^ with minor modifications. Cells from serum bottles and chemostats were mixed with BSA-treated 10 nm colloidal gold solution and 4 μl of this mixture were applied to a glow-discharged, X-thick carbon-coated, R2/2, 200 mesh copper Quantifoil grid (Quantifoil Micro Tools) in a Vitrobot chamber (FEI). Excess liquid was blotted off with a blot force of 6, blot time of 3 s, and drain time of 1 s and the grid was plunge-frozen for ECT imaging. All perfusion samples were on glow-discharged, X-thick carbon-coated, R2/2, Au NH2 London finder Quantifoil EM grids (Quantifoil Micro Tools) and were blotted either manually or automatically using the Vitrobot after addition of 1.5 μl gold fiducial markers. Imaging of all samples was performed on an FEI Polara 300-keV field emission gun electron microscope equipped with a Gatan image filter and K2 Summit counting electron-detector camera (Gatan). Data were collected using the UCSFtomo software^59^, with each tilt series ranging from −60° to 60° in 1° increments, an underfocus of ~5-10 µm, and a cumulative electron dose of ~130-160 e/A^2^ for each individual tilt series. The IMOD software package was used to calculate 3D reconstructions^60^.

### ECT Data Analysis and structure overlay of MtrC and MtrA on EM map

For figure (6C) and Movie S11 densities on the membrane interior and exterior were labeled with model points manually using the IMOD software. Density coordinates were extracted and used for distance calculations. The number of model points in (6C) is greater than the densities observed in (6A) since (6C) represents all the densities seen in the entire tomogram and (6A) is a slice from the tomogram that does not contain all the densities. For Figure 6B, crystal structure of MtrC was visualized and oriented using UCSF Chimera^61^. Adobe Photoshop was then used to overlay this crystal structure onto densities on the EM map. Surface view of MtrA was visualized using the PyMOL software^62^ and overlaid on EM densities using Adobe Photoshop.

### Direct Tunneling Limit Calculation

To find whether direct tunneling is possible between the observed densities in Fig. 6A, we used the available structures of MtrC and MtrA to calculate the largest center-to-center intermolecular distance that would allow direct tunneling. From the crystal structure, the dimensions of MtrC are ~9×6×4 nm^16^. Since the orientation of MtrC on the outer membrane is unknown, we took the largest dimension (9 nm) to calculate the direct tunneling limit. The known direct tunneling limit for the distance between the donor and acceptor redox sites in biological ET is ~2 nm^1^. This sets the limit of direct tunneling between MtrCs at 11 nm (9 nm + 2 nm) center-to-center intermolecular distance, assuming redox sites are located at the edges of the molecule. In the case of MtrA, the known molecular dimensions are ~10×5×2.5 nm^30^. From the EM map, it appears that MtrA is oriented with its long axis (10 nm) perpendicular to the outer membrane (Fig. 6B). Therefore, it is likely that any intermolecular electron tunneling between MtrAs occurs along the shorter axes (5 nm and 2.5 nm). Taking the larger of these two dimensions (5 nm), we calculate the limit of direct tunneling between MtrAs to be at 7 nm (5 nm + 2 nm) center-to-center intermolecular distance.

### Diffusion Timescale Calculation

To calculate the timescale of diffusion-based electron transfer events, we consider diffusion-based collisions between cytochromes that are diffusing in the membrane. We calculate the timescale of diffusion-based collision between the two reacting particles in a two dimensional membrane using the Hardt approach^63^:

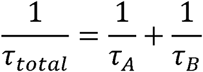

Here, *τ_total_* is the mean transit time for the reaction, is the mean time it takes a diffusing A particle to find an immobilized B particle and is the mean time it takes a diffusing B particle to find an immobilized A particle. *τ_A_* and *τ_B_*can be calculated from the diffusion transit time expression in two dimensions^63,64^: 

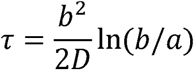

where *b* is the diffusion distance, *a* is the center-to-center distance between the reacting particles when reaction (in our case, electron transfer) occurs, and *D* is the diffusion coefficient of the diffusing particle. In cytochrome-cytochrome reactions, *a* was taken to be the average size of the cytochrome (6.33 nm for MtrC/OmcA, 3.75 nm for MtrA), *b* was taken to be the average nearest-neighbor distance found in the tomogram (Fig. 6I) (15.5 nm for putative MtrC/OmcA, 11.2 nm for putative MtrA), and *D* was assumed to be ~3 µm^2^/s for both MtrC/OmcA and MtrA^57^. However, in finding the average, we excluded all the nearest-neighbor distances that lay within the direct tunneling limit (< 11 nm for MtrC/OmcA, < 7 nm for MtrA). Finally, using the above values, we calculated *τ_total_* in Equation 1 for MtrC/OmcA to be 1.8×10^-5^ s and for MtrA to be 1.1×10^-5^ s.

